# Concurrent inhibition of ICMT and RAF/MEK suppresses RAC1^P29S^-driven MAPKi resistance in BRAF^V600E^ melanoma by regulating TAZ activity

**DOI:** 10.1101/2025.08.10.669587

**Authors:** Xiaoyang Gu, Patrick J Casey, Mei Wang

## Abstract

The RAC1 GTPase hotspot mutation P29S (RAC1^P29S^) is among the top driver oncogenes of cutaneous melanoma, which is known to develop resistance to MAPK pathway inhibitors including those targeting BRAF and MEK. Isoprenylcysteine carboxyl-methyltransferase (ICMT) is the enzyme catalyzing the last step of post-translational prenylation of RAC1, which is among its substrates. We demonstrate that RAC1^P29S/C189S^, which lacks C-terminal prenylation site, has lost the ability to induce resistance toward vemurafenib or trametinib in BRAF^V600E^ melanoma cells. Furthermore, the combination of vemurafenib with cysmethynil, a proof-of-concept ICMT inhibitor, showed efficacy in combating RAC1^P29S^-driven resistance of BRAF^V600E^ melanoma cells in both *in vitro* and *in vivo* settings. Concurrent treatment with cysmethynil and the MAPK pathway inhibitors efficiently inhibited proliferation and tumor formation of RAC1^P29S^ cells that are resistant to MAPK pathway inhibitor alone. Mechanistically, we found that the combined treatment impaired the nuclear translocation of TAZ, whose transcriptional activity is shown to account for MAPKi resistance in RAC1^P29S^ melanoma. We further validated the role of TAZ in RAC1^P29S^-driven resistance by demonstrating that introducing a constitutively-active TAZ mutant enhanced the MAPKi resistance in native cells, phenocopying the effect of RAC1^P29S^. The novel application of MAPKi and cysmethynil combination in RAC1^P29S^-driven MAPKi-resistant melanoma cells extends the potential utility of ICMT inhibitors, and also provides a new mechanism for targeting ICMT in cancer.

## Introduction

The incidence of cutaneous melanoma continues to increase, especially that in association with ultraviolet exposure (1,2). Although the prognosis of early-stage cutaneous melanoma has been improved over the past decade, the 5-year relative survival of distant metastatic melanoma is still quite poor and fraught with drug resistance (3). The top two driver oncogenes in melanoma are BRAF and NRAS, which are mutated in 63% and 26% of the cases, respectively (4), with 70-88% of BRAF mutations being V600E (5–7). The combination of BRAF and MEK inhibitors is one of the standard treatment regimens for BRAF-mutated melanoma, but despite good responses in the beginning all three FDA-approved combinations (vemurafenib + cobimetinib, dabrafenib + trametinib, and encorafenib + binimetinib) develop acquired resistance in about 50% patients within 12 months (2,8–14).

RAC1 is a small GTPase of RHO family. Although the major alteration of RAC1 in cancer is gene amplification, it is reported to be often mutated in melanoma with an overall rate around 4-8% (15). RAC1^P29S^ is the most common RAC1 mutation and the 3^rd^ most frequent activating mutation, next only to BRAF and NRAS in a UV associated-cohort (4,16). The RAC1^P29S^ mutant has higher GTP-bound level and exists mostly as activated form (4,17). While the association of RAC1^P29S^ melanoma patients with early resistance to RAF and other MAPK pathway inhibitors was reported a decade ago (18,19), it has been an ongoing challenge to find therapeutics options to counter the resistance.

Prenylation-dependent processing constitutes three-step post-translational modifications for substrate proteins, with the final step exclusively catalyzed by the enzyme isoprenylcysteine carboxylmethyltransferase (ICMT) (20). Processing by ICMT is critical for the functions of its substrates. It has been reported that ICMT knockout can abrogate membrane localization of KRAS (21) and abolish tumorigenesis driven by mutants of all RAS isoforms (22), supporting the potential of targeting ICMT in cancer. Cysmethynil is a proof-of-concept inhibitor that has been evaluated on several cancer cell lines and xenograft models and found to be effective in inducing autophagic cell death and apoptosis (22–25). Besides RAS, RAC1 is a prototypical ICMT substrate whose function can be affected by the prenylation processing. In this study, we evaluated the impact of ICMT inhibitor on the drug sensitivity to MAPK pathway inhibitors of RAC1^P29S^-BRAF^V600E^ melanoma cancer cells.

## Results

### Introducing Active RAC1 mutant protein in BRAF^V600E^ melanoma cells lead to MAPKi resistance

In cutaneous skin melanoma, missense RAC1 constitutes the third most frequent driver-mutations (26–29), most of which are P29S and P29L. Interestingly, RAC1 mutations are often present with BRAF or NRAS mutations (Figure S1A-B). Consistently, a recent study has shown that 47% and 28% of RAC1-mutant samples contain NRAS or BRAF mutations, respectively (30). Moreover, RAC1 mutation significantly reduces disease free survival independent of BRAF or NRAS status, and the presence of double mutants of RAC1-BRAF or RAC1-NRAS results in worse survival than those of any single mutation (Figure S1C).

To study the consequences of introducing various RAC1 proteins on treatment sensitivity of BRAF^V600E^ melanoma cancers, we introduced RAC1 variants, including RAC1^P29S^, RAC1^Q61L^ (GTP hydrolysis deficient), RAC1^WT^, and RAC1B (RAC1 splicing variant) (31) into BRAF^V600E^ melanoma A375 and A101D cells (supplementary Figure S2A-B). These cells were then subjected to the treatment by vemurafenib (BRAF^V600E^ inhibitor) or trametinib (MEK inhibitor). We found that the expression of the active RAC1 mutants - RAC1^P29S^ and RAC1^Q61L^ – led to higher tolerance to vemurafenib or trametinib both under the 2D-adherence culturing condition (Figure 1A-D) and in anchorage–independent soft agar colony formation assays (Figure 1E-H and supplementary Figure S2C-F). In contrast, RAC1^WT^ and RAC1B did not confer the cells with resistance to vemurafenib or trametinib (Figure 1E-H; the complete growth curves of soft agar assay are shown in Supplementary Figure S2C-F), suggesting that it is the RAC1 activity, not the level of protein expression, that is accountable for the drug resistance. To support this conclusion, we quantified the level of expression for both the exogenously introduced RAC1 isoforms and the total RAC1 by PCR (Figure S2A-B), which confirmed that the level of expression among the isoforms is not the reason for resistance from the two active forms of RAC1.

**Figure 1.**
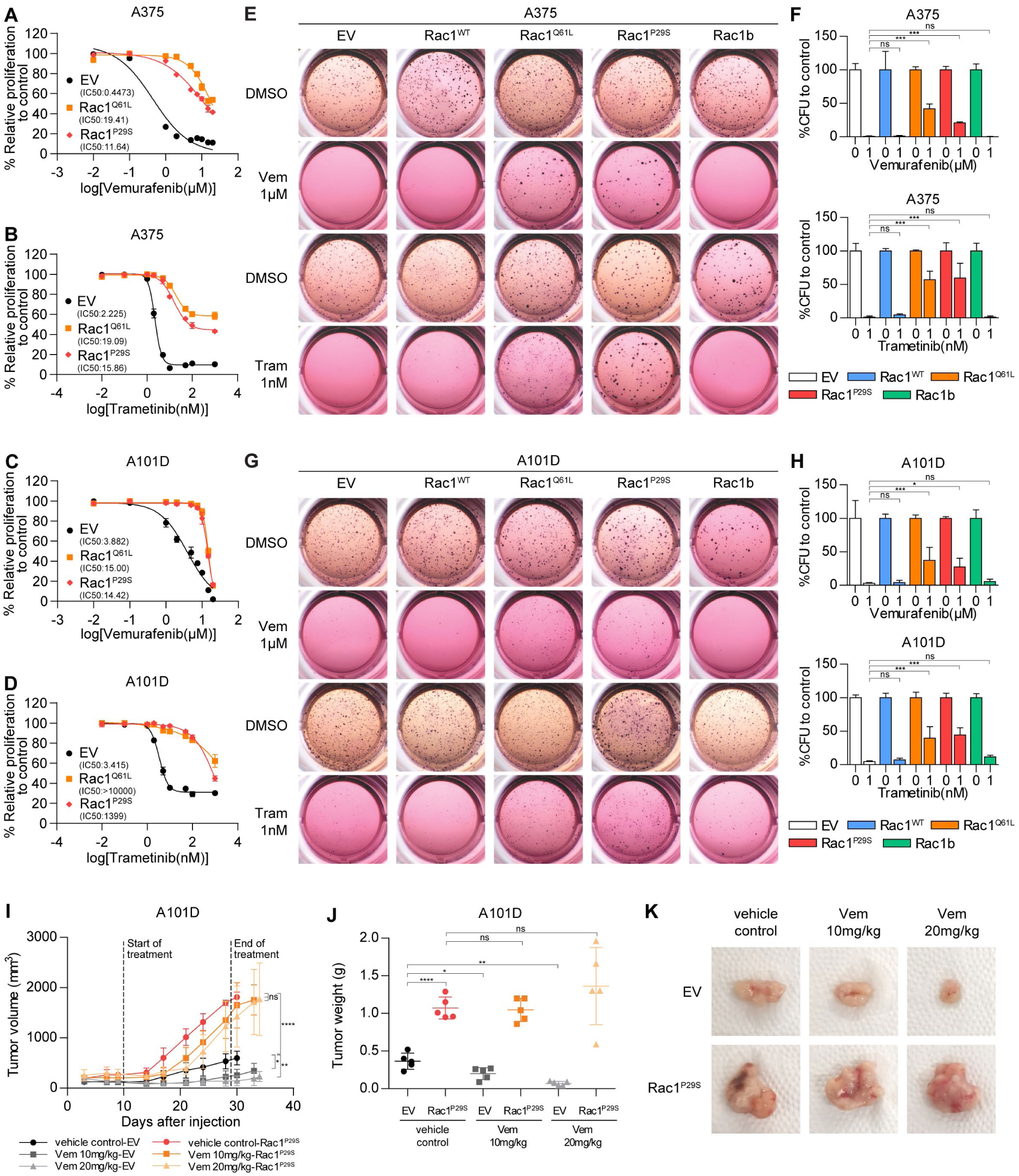
Introducing constitutively active RAC1 mutant proteins in BRAF^V600E^ melanoma cells leads to MAPKi resistance (A-B) Proliferation inhibition curves of A375 cells transduced with empty vector (EV), RAC1^Q61L^, or RAC1^P29S^ after 72h treatment of (A) vemurafenib ranging from 0.01µM to 20µM, and (B) trametinib ranging from 0.01nM to 1000nM. **(C-D)** Proliferation inhibition curves of A101D cells transduced with EV, RAC1^Q61L^, or RAC1^P29S^ after 96h treatment of (C) vemurafenib ranging from 0.01µM to 20µM, and (D) trametinib ranging from 0.01nM to 1000nM. **(E)** Representative soft agar colony images of A375 cells expressing EV, RAC1^WT^, RAC1^Q61L^, RAC1^P29S^, or RAC1B after 14-day incubation with 1µM vemurafenib or 1nM trametinib; comparisons are made with DMSO. **(F)** Normalized colony numbers of replicates from the experiment of (E). Statistical significance was determined by one-way ANOVA with Dunnett’s test: ns, not significant; ***p < 0.001. **(G)** Representative soft agar colony images of A101D cells expressing EV, RAC1^WT^, RAC1^Q61L^, RAC1^P29S^, or RAC1B after 14-day incubation with 1µM vemurafenib or 1nM trametinib; comparisons are made with DMSO. **(H)** Normalized colony numbers of replicates from the experiment of (G). Statistical analysis as described in (F): *p < 0.05, ***p < 0.001. **(I)** Growth curves of xenograft tumors derived from A101D cells transduced with EV or RAC1^P29S^ under b.i.d. oral treatment with vehicle, 10mg/kg or 20mg/kg vemurafenib; n = 5 for each dosing group. For each mouse, the EV and RAC1^P29S^ cells were implanted on the contralateral side on Day 0; treatment was started on Day 10 and ended on Day 29. Statistical significance was determined by two-tailed unpaired t-test with Welch’s correction: ns, not significant; *p < 0.05, **p < 0.01, ****p < 0.0001. **(J)** Endpoint tumor weights are plotted for each group from (I). End time point for each group was determined by IACUC protocol to be one of the two tumor in the same mouse reached the size of 2000 mm^3^: vehicle control, Day 30; 10mg/kg vemurafenib, Day 33; 10mg/kg or 20mg/kg vemurafenib, Day 34. Mean tumor weight ± S.D. is presented: EV vehicle control, 0.366 ± 0.107g; RAC1^P29S^ vehicle control, 1.072 ± 0.146g; EV Vem 10mg/kg, 0.202 ± 0.079g; RAC1^P29S^ Vem 10mg/kg, 1.046 ± 0.155g; EV Vem 20mg/kg, 0.070 ± 0.028g; RAC1^P29S^ Vem 20mg/kg, 1.362 ± 0.512g. Statistical significance was determined by two-tailed unpaired t-test with Welch’s correction: ns, not significant; *p < 0.05, **p < 0.01, ****p < 0.0001. **(K)** Representative images for tumors in (J) at sacrifice.

The evaluation of the impact of RAC1^P29S^ was extended to the *in vivo* xenograft model by subcutaneously implanting the A101D cells stably expressing RAC1^P29S^ or empty vector control into the contralateral sides or of NOD SCID mice. Consistent with the *in vitro* findings, RAC1^P29S^-expressing A101D xenograft sustained faster growth under vemurafenib treatment, compared to the control tumors (Figure 1I-K and Figure S3A). The dosages of vemurafenib were well-tolerated as the body weights of all treatment groups showed no significant difference (Figure S3B).

### RAC1^P29S^-driven malignant transformation and MAPKi resistance depend on its C-terminal post-translational modification

It is well-known that mutant RAS proteins are driver mutations for malignant transformation and maintenance. Despite the frequent presence of RAC1 activation mutations in cancers, the direct link of RAC1 in transformation is not as well-recognized. Having shown the RAC1 activation leads to MAPKi resistance in BRAF mutant melanoma cells, we confirmed the transforming ability of RAC1^P29S^, and further, evaluated the importance of its C-terminal modification for this functional ability. To test transforming capability, the same RAC1 isoforms as described above were expressed in MCF10A (Figure S4A), a benign human mammary epithelial cell line. Both RAC1^Q61L^ and RAC1^P29S^ expression enabled anchorage-independent colony formation – a characteristic of malignant cells – in MCF10A cells, while the control cells and those expressing RAC1^WT^ or RAC1B failed to form colonies (Figure 2A). This evidence supports the role of constitutively-active RAC1 as an oncogenic driver that can independently transform cells. Given that RAC1 is a well-known prenylation processing substrate, we evaluated the functional impact of ICMT on RAC1^P29S^-driven transformation. We found that treatment with the ICMT inhibitor, cysmethynil, effectively abolished colony formation of RAC1^P29S^-expressing MCF10A cells (Figure 2B). RAC1 GTPase activity assay confirmed that cysmethynil treatment reduced the level of GTP-bound RAC1 in A101D cells (Figure 2C). Taken together, the results demonstrate the potential of ICMT inhibition in RAC1-driven cancers.

**Figure 2.**
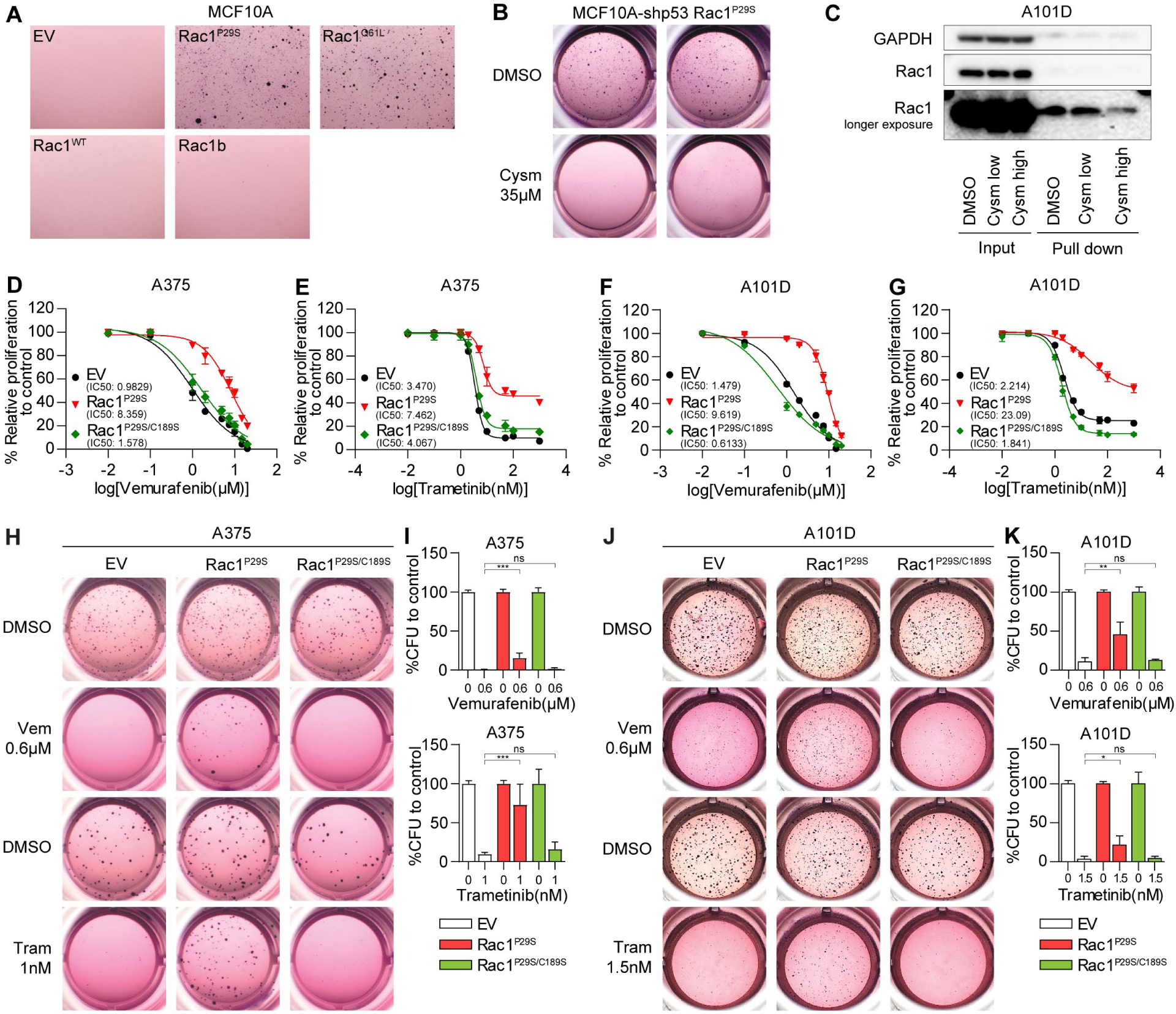
RAC1^P29S^-driven malignant transformation and MAPKi resistance depend on its C-terminus post-translational modification. **(A)** Representative images of soft agar colony formation assay of MCF10A cells transduced with EV, RAC1^WT^, RAC1^Q61L^, RAC1^P29S^, or RAC1B after 14-day incubation. **(B)** Soft agar colony formation of MCF10A cells transformed by expressing of p53 shRNA and RAC1^P29S^ after 14-day treatment of DMSO or 35µM cysmethynil. **(C)** RAC1 activity assay by pulldown of GTP-bound RAC1 on A101D cells stably expressing RAC1^P29S^ after 48h treatment by DMSO or cysmethynil. **(D-E)** Proliferation inhibition curves of A375 cells transduced with EV, RAC1^P29S^, or RAC1^P29S/C189S^ subjected to 72h treatment of (D) vemurafenib ranging from 0.01µM to 20µM, and (E) trametinib ranging from 0.01nM to 1000nM. **(F-G)** Proliferation inhibition curves of A101D cells transduced with EV, RAC1^P29S^, or RAC1^P29S/C189S^ subjected to 96h treatment of the same drug treatment as in (D-E). **(H)** Representative colony images of A375 cells expressing EV, RAC1^P29S^, or RAC1^P29S/C189S^ after 14-day incubation with 0.6µM vemurafenib or 1nM trametinib. **(I)** Normalized colony numbers of all replicates from experiment of (H). Statistical significance was determined by one-way ANOVA with Dunnett’s test: ns, not significant; ***p < 0.001. **(J)** Representative colony images of A101D cells expressing EV, RAC1^P29S^, or RAC1^P29S/C189S^ after 14-day incubation with 0.6µM vemurafenib or 1.5nM trametinib. **(K)** Same analysis as described in (I) on all replicates from experiment of (J): *p < 0.05, **p < 0.01.

In addition to pharmacologic inhibition, tumor-promoting functions were compared between RAC1^P29S^ and RAC1^P29S/C189S^ – the mutant that lacks C-terminal prenylation modification. RAC1^P29S^ and RAC1^P29S/C189S^ were introduced into A375 and A101D melanoma cells together with empty vector control (Figure S4B-C). Unlike RAC1^P29S^, expression of RAC1^P29S/C189S^ did not confer resistance to vemurafenib and trametinib under adherent culture (Figure 2D-G) and soft agar anchorage independent growth condition (Figure 2H-K; the complete growth curves of soft agar assay are shown in Figure S4D-G), supporting the conclusion from ICMT inhibitor treatment that C-terminal modification is indispensable for RAC1^P29S^-driven resistance to MAPKi.

### ICMTi and MAPKi work synergistically to combat RAC1^P29S^-driven resistance in both *in vitro* and *in vivo* settings

Having confirmed the importance of RAC1 C-terminal modification for its function in driving tumorigenesis, we evaluated the impact on overcoming the RAC1^P29S^-driven MAPKi resistance by adding ICMTi – cysmethynil - to BRAFi or MEKi in the treatment. In RAC1^P29S^-expressing A375 and A101D cells, the combination treatment elicited responses that MAPKi alone could not induce (Figure S5A-D). Comparing parental A375 and A101D cells with those expressing RAC1^P29S^, we found the combination treatment effectively inhibited the proliferation of the RAC1^P29S^-expressing cells to the similar level of that of parental cells in response to single MAPKi – vemurafenib or trametinib (Figure 3A-D). The dose-response matrices of vemurafenib + cysmethynil and trametinib + cysmethynil combinations were determined to evaluate the drug-drug interactions. Synergy scores calculated with Bliss independence model and plotted by SynergyFinder+ (32,33) demonstrate that both combinations were synergistic in suppressing proliferation of A375 and A101D cells expressing RAC1^P29S^, especially at higher dosages (Figure 3E-H).

**Figure 3.**
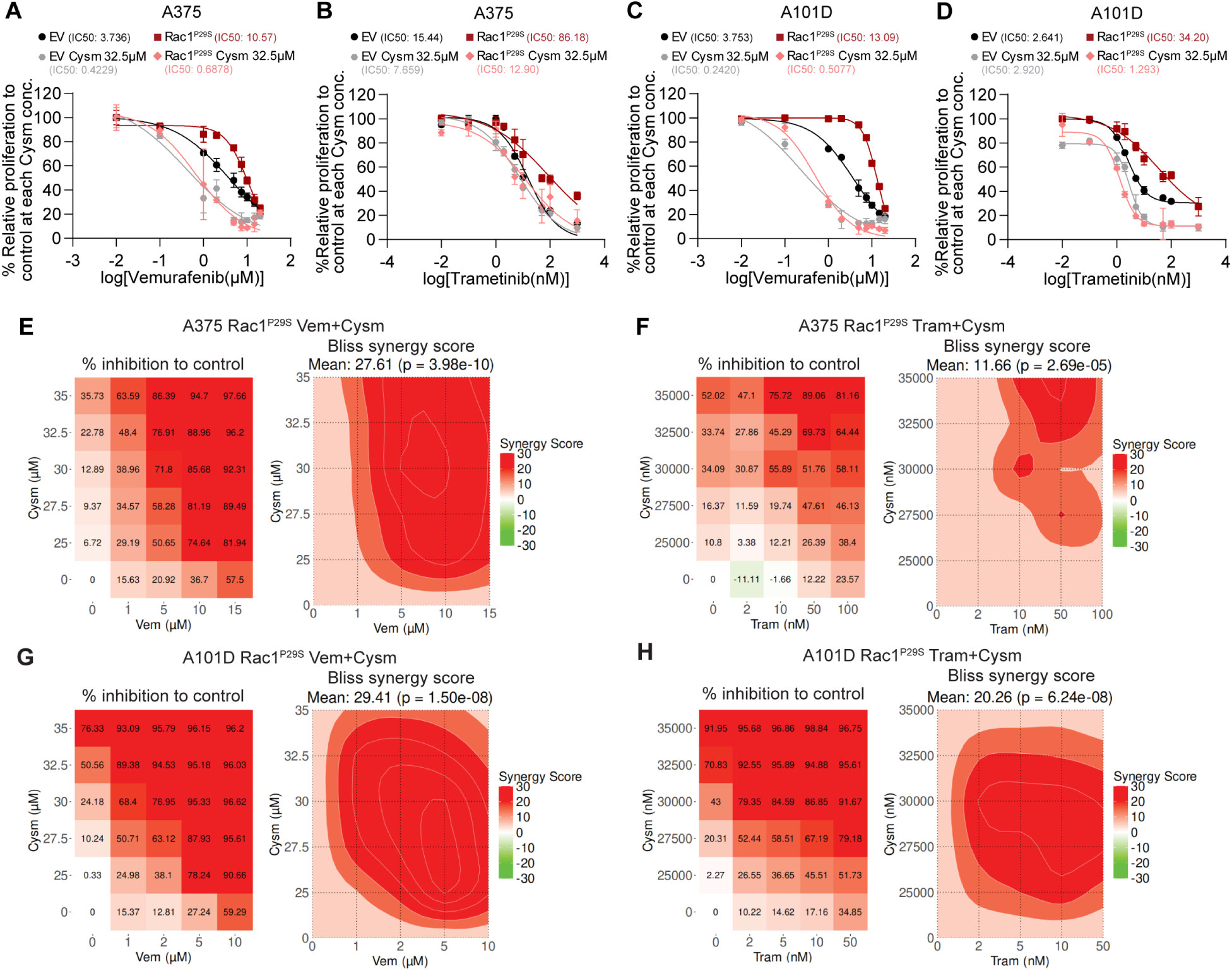
Combination of MAPKi and ICMTi displays synergistic anti-proliferative effect on BRAF^V600E^ + RAC1^P29S^ melanoma cells. (A-B) Proliferation inhibition curves of A375 cells expressing EV or RAC1^P29S^ subjected to 72h treatment of (A) 0.01-20µM vemurafenib with or without 32.5µM cysmethynil, and (B) 0.01-1000nM trametinib with or without 32.5µM cysmethynil. **(C-D)** Proliferation inhibition curves of A101D cells expressing EV or RAC1^P29S^ subjected to 72h treatment of (C) 0.01-20µM vemurafenib with or without 32.5µM cysmethynil, and (D) 0.01-1000nM trametinib with or without 32.5µM cysmethynil. **(E-F)** Inhibition plots and Bliss synergy score maps of A375 cells transduced with RAC1^P29S^ after 72h concurrent treatment by (E) 1-15µM vemurafenib plus 25-35µM cysmethynil, and (F) 2-100nM trametinib plus 25-35µM cysmethynil. **(G-H)** Inhibition plots and Bliss synergy score maps of A101D cells transduced with RAC1^P29S^ after 72h concurrent treatment by (G) 1-10µM vemurafenib plus 25-35µM cysmethynil, and (H) 2-50nM trametinib plus 25-35µM cysmethynil.

Evaluation of the combination treatment was extended to the anchorage-independent growth condition with soft agar colony formation assays. Not surprisingly, the cysmethynil and MAPKi combination synergistically reduced colony formation of RAC1^P29S^-expressing A375 and A101D cells (Figure 4A-D). Lastly, we evaluated the effect of combined treatment in the xenograft mouse model. The cysmethynil (150mg/kg/48h) and vemurafenib (20mg/kg/b.i.d) treatment regimens follow the published methods (23,25). Consistent with the *in vitro* findings, the response of RAC1^P29S^-expressing A101D cell xenografts to vemurafenib was not significantly different from that of vehicle control treatment, whereas addition of ICMTi significantly enhanced the anti-proliferative efficacy of vemurafenib (Figure 4E-F). All treatment regimens were well tolerated as demonstrated by stable body weight (Figure 4G). Considering these *in vitro* and *in vivo* evaluations, we conclude that the synergistic effects are consistent with the notion that the ICMT inhibition can revert RAC1^P29S^-driven MAPKi resistance.

**Figure 4.**
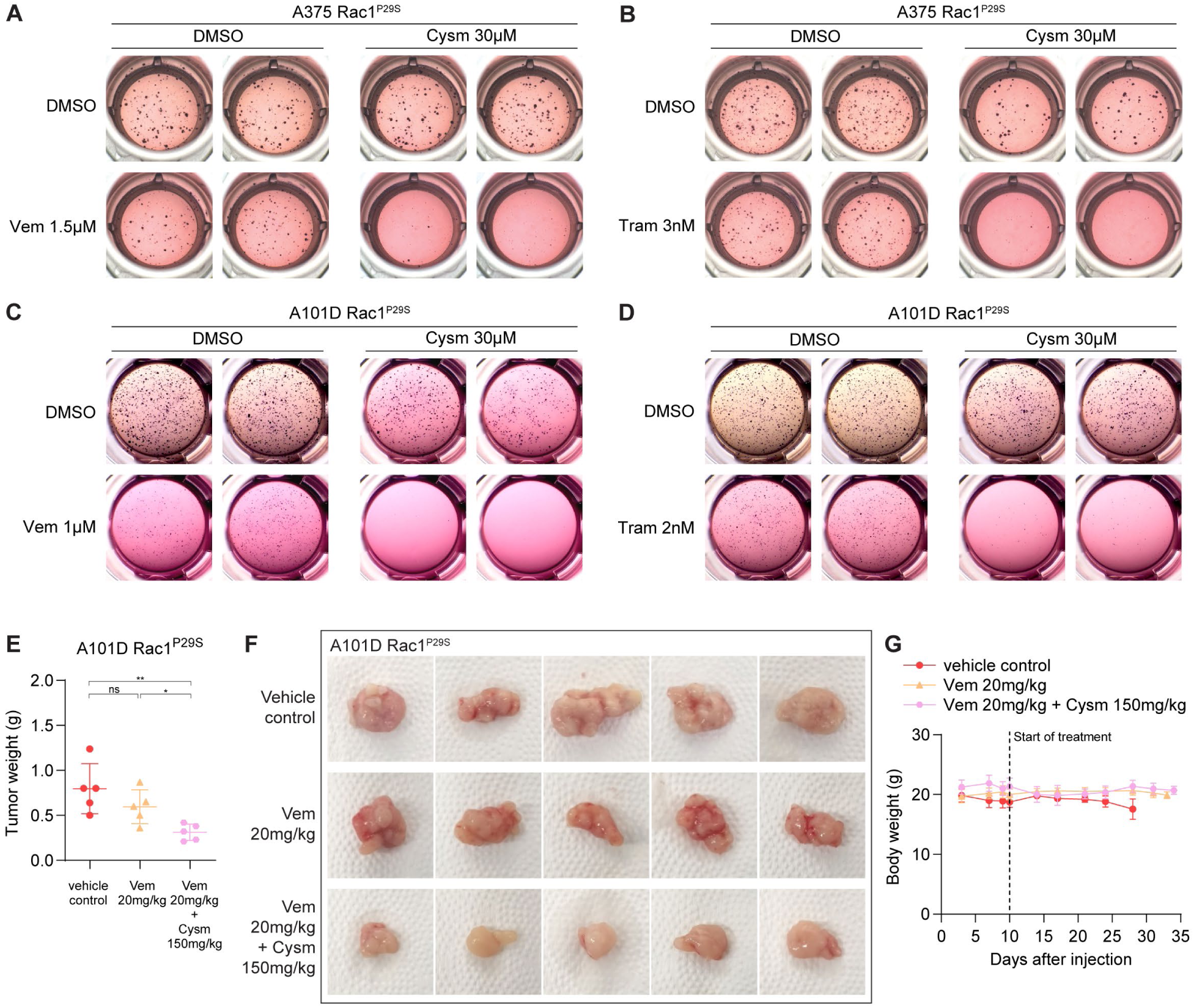
RAC1^P29S^-induced resistance to MAPKi can be obliterated by concurrent ICMT inhibition both under anchorage-independent growth condition and *in vivo* tumor growth. (A-B) Soft agar colony formation of A375 cells transduced with RAC1^P29S^ after 14-day incubation with (A) 1.5µM vemurafenib with or without cysmethynil, and (B) 3nM trametinib with or without cysmethynil. **(C-D)** Soft agar colony formation of A101D cells transduced with RAC1^P29S^ after 14-day incubation with (C) 1µM vemurafenib with or without cysmethynil, and (D) 2nM trametinib with or without cysmethynil. **(E)** Endpoint tumor weight for A101D-RAC1^P29S^ xenografts under twice a day oral dosing of vehicle or 20mg/kg vemurafenib with or without every other day treatment by 150mg/kg cysmethynil; n = 5 for each dosing group. Endpoint was determined by the tumor size reached 1000 mm^3^. Statistical significance of Mean tumor weight ± S.D. was determined by two-tailed unpaired t-test with Welch’s correction: ns, not significant; *p < 0.05, ***p < 0.001. **(F)** Images of tumors from (E) at the time of sacrifice. **(G)** Body weights for mice of each treatment group as described in Figure 4E.

### Combination treatment by MAPKi and ICMTi reduces nuclear TAZ level and TAZ-driven transcription

Although good efficacy has been observed with the combination of MAPKi and ICMTi, the molecular mechanism underneath the synergistic effects is not clear, particularly due to the complexity of cell signaling pathways involved and the cross-talks among the downstream proteins. In recent studies, we have found that ICMT regulates the cellular levels of TAZ but not YAP. Further, in KRAS-driven cancers, ICMT inhibition of tumorigenesis, at least partly, is mediated through its control of TAZ activity (34). Among the canonical ICMT substrates, RAC1 is a major regulator of actin-cytoskeleton organization, which is known to play a role in YAP/TAZ nuclear localization and could potentially confer MAPKi resistance (35,36). Interestingly, in these melanoma cell lines, we found that TAZ is dominantly expressed among the TAZ/YAP homologs. Hence, we evaluated the impact of vemurafenib, cysmethynil and their combination on nuclear and cytosolic levels of TAZ. Remarkably, we found that vemurafenib treatment effectively reduced nucleus TAZ in A375 and A101D control cells while having a minimal effect on those the expressing RAC1^P29S^, demonstrating a RAC1^P29S^-driven resistance to MAPKi control of TAZ localization (Figure 5A-B). Consistent with the hypothesis of TAZ playing a critical role in the efficacy of ICMTi and MAPKi, the combination treatment was able to reduce TAZ nuclear localization in the RAC1^P29S^-expressing cells (Figure 5A-B). To evaluate whether the TAZ localization response to the combination treatment is due to the diminished RAC1 C-terminal modification, we performed the same vemurafenib single agent treatment on RAC1^P29S^-and RAC1^P29S/C189S^-expressing A375 and A101D cells. Control cells and RAC1^P29S^ cells responded to vemurafenib treatment the same way as in Figure 5A-B, with RAC1^P29S^ cells resisting the vemurafenib regulated nuclear TAZ reduction (Figure 5C-D). RAC1^P29S/C189S^, instead, behaves similarly as when RAC1^P29S^ cells were treated by the ICMTi and vemurafenib combination, with a diminished nuclear TAZ level (Figure 5C-D), which suggests that active RAC1 can only promote TAZ nuclear localization when it has intact C-terminus prenylation processing.

**Figure 5.**
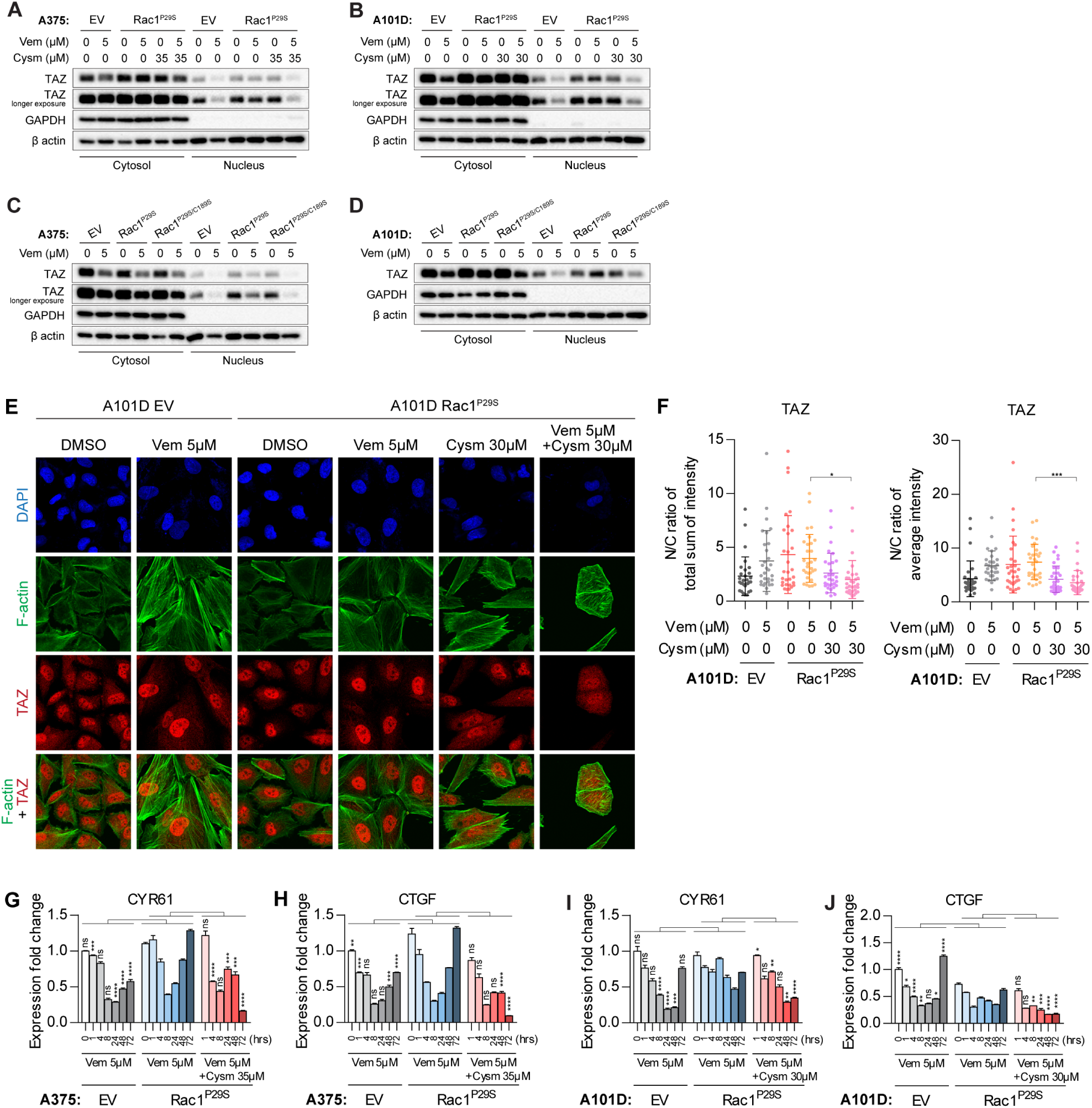
Combination treatment by MAPKi and ICMTi reduces nuclear TAZ level and TAZ-driven transcription in BRAF^V600E^/RAC1^P29S^ melanoma cells. **(A)** Immunoblot of indicated proteins on the cytosolic and nucleus fractions from A375 cells transduced with EV or RAC1^P29S^ after 48h treatment of 5µM vemurafenib with or without 35µM cysmethynil. **(B)** The same evaluation as in (A) was performed on A101D cells; the cysmethynil concentration is 30µM. **(C)** Immunoblot of indicated proteins on the cytosolic and nucleus fractions from in A375 cells expressing EV, RAC1^P29S^, or RAC1^P29S/C189S^ after 48h treatment of 5µM vemurafenib. **(D)** The same evaluation as in (C) was performed on A101D cells **(E)** Confocal microscope images for TAZ in A101D cells expressing EV or RAC1^P29S^ after 72h treatment of 5µM vemurafenib with or without 30µM cysmethynil. **(F)** Statistical analysis for nuclear and cytosol abundance of TAZ in A101D cells transduced with EV or RAC1^P29S^ on (E). 30 cells were analyzed by line plot profile and calculated for N/C ratio for each condition. Statistical significance was determined by one-way ANOVA with Tukey’s test: ns, not significant; *p < 0.05, ***p < 0.001. **(G-H)** Time-course qPCR analysis for TAZ regulated expression of (G) CYR61 and (H) CTGF in A375 cells expressing EV or RAC1^P29S^ after treatment of 5µM vemurafenib with and without concurrent cysmethynil treatment. Data is presented as mean ± S.E.M. Statistical significance was determined by two-way ANOVA with Bonferroni test (compared with RAC1^P29S^ Vem 5µM under each timepoint; RAC1^P29S^ Vem 5µM + Cysm 35µM 0h was defined to be the same as RAC1^P29S^ Vem 5µM 0h): ns, not significant; **p < 0.01, ***p < 0.001, ****p < 0.0001. **(I-J)** Time-course qPCR analysis for TAZ regulated expression of (I) CYR61 and (J) CTGF in A101D cells expressing EV or RAC1^P29S^ after same treatment and data analysis as in (G-H); the cysmethynil concentration is 30µM.

Providing further evidence for TAZ localization changes in response to RAC1^P29S^ and to the inhibitor treatment, we visualized TAZ by confocal immunofluorescence of A101D and A101D-RAC1^P29S^ cells under the same treatment as in Figure 5B. Consistent with the fractionation study, the quantitative analysis of the confocal images demonstrates that the nuclear TAZ level did not reduce in response to vemurafenib single agent treatment (in contrast to A101D-EV control cells), whereas the combination of ICMTi and vemurafenib significantly reduced nuclear TAZ level in these A101D-RAC1^P29S^ cells (Figure 5E-F).

TAZ acts as a transcription activator that forms a complex with transcription factor TEAD to regulate the expression of target genes. CYR61 (CCN1) and CTGF (CCN2), important regulators of tumorigenesis and proliferation (37,38), are canonical targets of TAZ-TEAD (39,40). To evaluate the consequences of TAZ localization in response to the treatments, we quantified the expression level of CYR61 and CTGF over several time points taking into account the time-dependent fluctuations of signaling pathways, especially the well-known MAPK related signals. This study revealed that vemurafenib did not change the pattern of CYR61 and CTGF expression, most notably the elevated expression in RAC1^P29S^ cells at late time points (Figure 5G-J). In contrast, the combination treatment of ICMTi and vemurafenib was able to suppress the expression of CYR61 and CTGF in RAC1^P29S^-expressing cells at late time points (Figure 5G-J), clearly demonstrated that the TAZ downstream targets changes are consistent with that of TAZ nuclear localization.

### TAZ is a major mediator for RAC1^P29S^-driven resistance to MAPKi in BRAF mutant melanoma

To evaluate the role of TAZ in RAC1^P29S^-driven MAPKi resistance, we performed TAZ knockdown (Figure S6A) study in A101D-RAC1^P29S^ cells. Consistent with the data so far, suppression of TAZ sensitized the A101D RAC1^P29S^ cells to vemurafenib and trametinib (Figure 6A-B). In parallel, the active TAZ^4SA^ (S66A, S89A, S117A and S311A) mutant was expressed in the parental A101D cells (Figure S6B), and the response to vemurafenib and trametinib was compared to that of A101D-RAC1^P29S^ cells. Consistent with the hypothesis based on the above results, TAZ^4SA^ expression, similar to RAC1^P29S^, led to MAPKi resistance (Figure 6C-D). Taken together, the evidence supports the role of RAC1, through TAZ-mediated gene expression, in the resistance of BRAF mutant melanoma cells to MAPKi, a common challenge for melanoma treatment. The results also provide rational for targeting ICMT in overcoming RAC1-activating-mutant-driven MAPK inhibitor resistance in not only melanoma but possibly in other cancers (Figure 6E).

**Figure 6.**
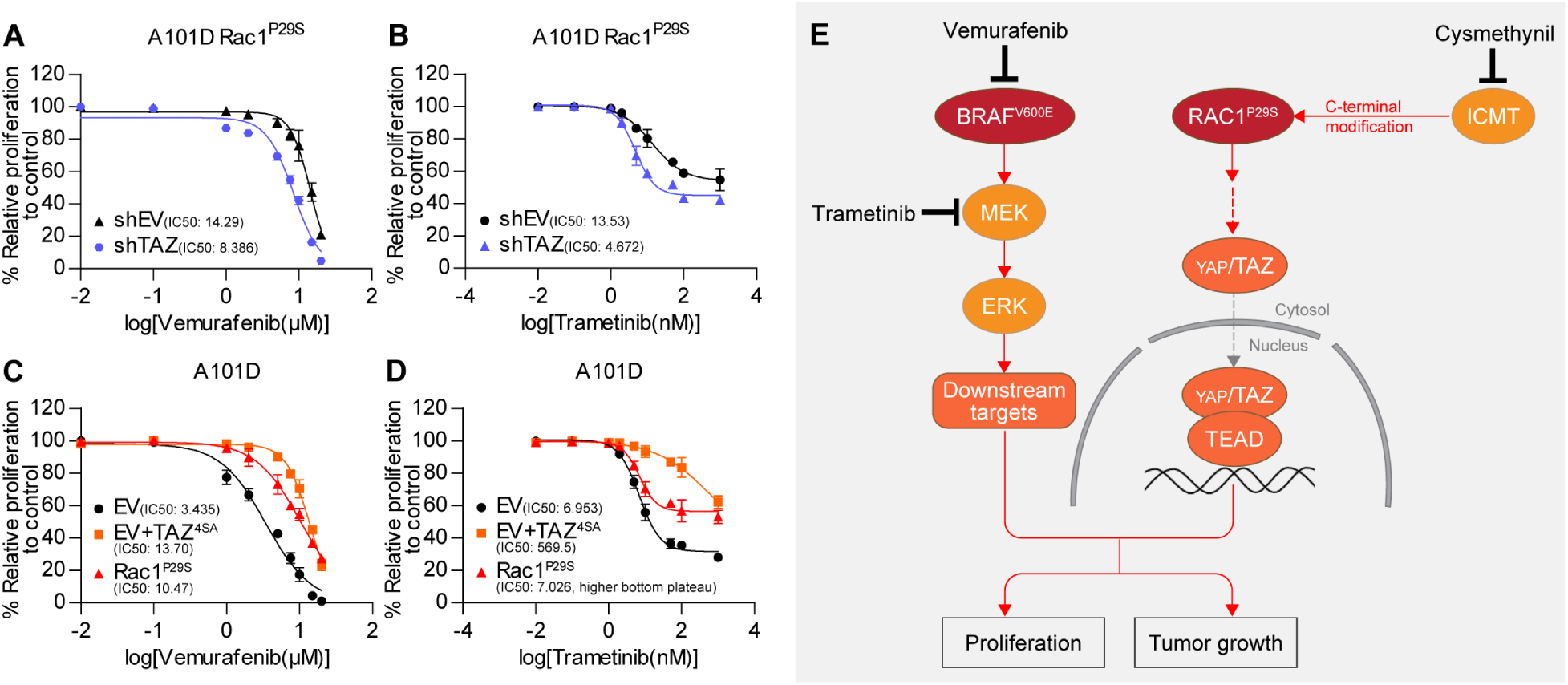
TAZ is a major mediator for RAC1^P29S^-driven resistance to MAPKi in BRAF mutant melanoma (A-B) Proliferation inhibition curves of A101D RAC1^P29S^ cells expressing TAZ-shRNA after 96h treatment of (A) vemurafenib ranging from 0.01µM to 20µM, and (B) trametinib ranging from 0.01nM to 1000nM. **(C-D)** Proliferation inhibition curves of A101D EV cells expressing TAZ^4SA^ compared with A101D-RAC1^P29S^ cells after 96h treatment of (C) vemurafenib ranging from 0.01µM to 20µM, and (D) trametinib ranging from 0.01nM to 1000nM. **(E)** Mechanism model for RAC1^P29S^-induced MAPKi resistance and the efficacy of MAPKi + ICMTi combination treatment. Briefly, the commonly used BRAF^V600E^ and MEK inhibitors will suppress melanoma progression through down-regulating the MAPK pathway that is over-activated by BRAF^V600E^. RAC1^P29S^ mutation drives MAPKi resistance in melanoma by enhancing TAZ nuclear localization and TAZ mediated gene transcription. As an ICMT substrate, RAC1^P29S^ function depends on its C-terminal modification. Hence, concurrent treatment by ICMTi and MAPKi overcome resistance and suppress proliferation both *in vitro* and *in vivo*. Red arrow: positive effects; black arrow: inhibitory effects; grey arrow: translocation event.

## Discussion

RAC1 GTPase is one of the so-called CAAX proteins, which contain Cysteine-Aliphatic-Aliphatic-Any amino acid sequences that serve as the consensus for prenylation processing at the C-termini. There are three enzymatic steps in the post-translational prenylation modification: it starts with the addition of either a 15-carbon or 20-carbon isoprenoid group to the cysteine residue, is followed by proteolysis to remove the –AAX peptide, and ends with carboxylmethylation of the cysteine by ICMT. These modifications are important for the transport, localization, stability, and protein–protein interactions of these substrate proteins, and thus for their proper function. Among the CAAX proteins, RAS proteins—particularly the KRAS isoform—garnered the most attention due to their role in tumorigenesis in a large number of human cancers. Prenylation processing gained attention during the era when RAS was considered “undruggable,” and targeting RAS modification was pursued as an alternative approach to inhibit its tumorigenic activity.

Although not as prominent in tumorigenesis as RAS proteins, RAC1 is increasingly recognized as a central signaling hub in cancer, involved in proliferation, survival, cytoskeletal dynamics, invasion, and microenvironment modulation. RAC1 also acts downstream of—and often in cooperation with—oncogenic RAS in several cancer pathways. While less frequently mutated than RAS, RAC1 hyperactivation is commonly achieved via overexpression, upstream drivers (including RAS), and activating mutations—dominated by the P29S mutation—which drive core cancer signaling and hallmark behaviours in both solid and hematologic malignancies. In addition to these critical roles, RAC1 activation is also recognized to promote therapy resistance, notably to RAS–MAPK-pathway inhibitors, which is the focus of this study. Despite the obvious value of targeting RAC1, it remains a significant challenge and an area of active exploration.

In our recent studies, we have found that inhibiting ICMT is effective in combating RAS-driven signaling pathways and therefore RAS-driven tumorigenesis (22,25,34). In this study, we have shown that the RAC1P29S/C189S mutant, lacking the C-terminal prenylation processing, lost its ability to promote proliferation and induce resistance to MAPKi, illustrating the potential of targeting prenylation processing in combating RAC1-driven cancer signaling. Reinforcing the conclusions from the RAC1P29S/C189S study, ICMT inhibitor treatment similarly abolished RAC1P29S-induced proliferation and MAPKi resistance. One of the major challenges for RAS– MAPK-pathway inhibitors, including those targeting RAS(G12C), is the development of adaptive resistance; hence, the results of the ICMTi combination study illustrate an attractive approach to combat resistance not only in RAC1-mutant cancers but potentially in other cancers, including those driven by mutant RAS. Notably, since our recent studies have shown that ICMT inhibition can suppress both RAS-driven (22,25,34) and RAC1-driven processes, there is significant potential for ICMT inhibitor treatment either as a single agent or in combination.

Mechanistically, we have shown that constitutively active RAC1 supports TAZ nuclear localization and transcriptional activity, which significantly contributes to MAPKi resistance. We speculate that RAC1’s function in regulating actin cytoskeleton organization plays a role in TAZ stability and localization. However, in-depth investigation will be required to elucidate the precise mechanism of this regulation.

## Materials and Methods

### Cell lines and culture conditions

All cell lines were purchased from ATCC and cultured according to ATCC instructions: A375 (CRL-1619), A101D (CRL-7815), MCF10A (CRL-10317), HEK293T (CRL-3216), MCF-7 (HTB-22). Cell proliferation was monitored in real-time using an IncuCyte S3 or ZOOM instrument (Sartorius) as previously described (41), with confluence quantified using IncuCyte ZOOM 2018A software. Soft agar colony formation assays and drug sensitivity determinations were performed as described in reference (34).

### Plasmids, vector cloning and site-specific mutagenesis

Plasmids: pBABE-puro (#1764, Addgene), pMSCV-Blasticidin (#75085, Addgene), pLL3.7-EGFP (#11795, Addgene); pLVX-puro (#632164, Takara Bio). All constructs were verified by Sanger sequencing. pBABE-zeo-RAC1-WT, pBABE-zeo-RAC1-Q61L, pMSCV-Blasticidin-flag-TAZ-WT and pMSCV-Blasticidin-flag-TAZ-4SA were previously generated in the lab. TAZ-4SA harbours S66A, S89A, S117A, S311A mutations. RAC1B isoform was PCR-amplified from MCF-7 cDNA and cloned into pLVX-puro. RAC1P29S was generated from RAC1WT using the QuikChange Site-Directed Mutagenesis Kit (#200518, Agilent) in pLVX-puro. RAC1^P29S/C189S^ was generated from RAC1^P29S^ by site-directed mutagenesis. pLL3.7-EGFP was used for shRNA cloning. Lentivirus and retrovirus production and transduction followed the reference (34).

### Chemicals, antibodies and reagents

Cysmethynil (2-[5-(3-Methylphenyl)-1-octyl-1H-indol-3-yl] acetamide) was synthesized by the Duke Small Molecule Synthesis Facility as previously described (23,42,43). Vemurafenib (V-2800) and trametinib (T-8123) were obtained from LC Laboratories. For western blot analysis, we used antibodies against GAPDH (#2118, Cell Signaling Technology - CST), YAP/TAZ (#8418, CST), and β-actin (AM1829B, Abgent), with detection using Goat anti-Rabbit and Goat anti-Mouse secondary antibodies (#31460 and #31430 from Thermo Fisher Scientific). For immunofluorescence, primary antibody against TAZ (HPA007415, Sigma-Aldrich) was detected with Alexa Fluor 568 Goat anti-Rabbit secondary antibody (A-11036, Invitrogen), with F-actin visualized using Alexa Fluor 488 Phalloidin (#8878, CST) and nuclei stained with DAPI (#62248, Thermo Fisher Scientific).

### Western blot and PCR analysis

Cells were seeded and allowed to adhere overnight before drug treatment. Cells were harvested, lysed, and proteins separated by SDS-PAGE, transferred to membranes, and probed with primary antibodies followed by HRP-conjugated secondary antibodies as described in reference (24). Chemiluminescent signals were detected. For subcellular fractionation, cell pellets were lysed hypotonically, suspended in NP-40 buffer, homogenized, and centrifuged (3,000 rpm) to separate cytosolic and nuclear fractions; fractions were then lysed and analyzed by immunoblotting per reference (44).

### RAC1 GTP-binding activity assay

RAC1-GTP levels were measured using the RAC1 Pull-down Activation Assay Biochem Kit (#BK035, Cytoskeleton Inc.) according to the manufacturer’s instructions and reference (41). Briefly, cells were seeded, attached overnight, treated as indicated, and processed per kit protocol. Bound RAC1-GTP was detected by immunoblotting.

### qRT-PCR analysis

Total RNA was extracted using the FavorPrep™ Tissue Total RNA Mini Kit (FATRK 001-2, Favorgen Biotech Corp) from treated cells. cDNA was synthesized using ReverTra Ace™ qPCR RT Master Mix (FSQ-201, Toyobo). qRT-PCR was performed as described before (34) using gene-specific primers and SYBR Green detection.

### Confocal immunofluorescence microscopy

Cells (8,000/well) were seeded in ibidi µ-Plates (#82406, ibidi) overnight, treated as indicated, fixed, permeabilized, blocked, and stained. Primary antibody (TAZ) incubation was overnight at 4°C. Secondary antibodies (Alexa Fluor 568), Phalloidin (Alexa Fluor 488), and DAPI were applied for 1 hour at room temperature. Samples were imaged on an Olympus FV3000 Confocal Microscope using a 40 x objective. Image acquisition settings were kept constant within experiments. Blinding and Randomization: Image acquisition order was randomized. Quantitative analysis of protein localization was performed using line plot profiles (ImageJ Plugin: RGB Profile Plot). Sample Size: 30 cells per condition, randomly selected from 3 independent experiments, were analyzed. Representative images are shown.

### Mouse tumor xenograft study

All procedures followed IACUC guidelines (Protocol # SHS/1462). Animals: Female NOD-SCID mice (NOD.CB17-Prkdcscid/J), aged 8-12 weeks, were used. A101D cells were mixed 1:1 (v/v) with Matrigel (#354234, Corning) and injected subcutaneously (5 x 10^5^ cells/injection, bilateral). Tumors were established for 10 days before treatment. Randomization: Mice were randomly assigned to treatment groups (n=8-10 tumors/group) after sizes reached ∼100mm³. Drug Administration: Vemurafenib (10 or 20 mg/kg) was dissolved in 5% DMSO/0.5% methyl cellulose (M0512, Sigma-Aldrich) and administered orally twice daily (50,51). Cysmethynil was prepared and administered intraperitoneally as described (23–25). Tumor measurements (caliper) and body weights were performed twice weekly. Tumor volume = (length x width²)/2.

## Statistical analysis

Data were plotted and analyzed using GraphPad Prism (v5.0). Numerical data are presented as mean ± SD unless otherwise stated. Specific statistical tests used are detailed in each figure legend: Kaplan-Meier survival curves (Log-rank (Mantel-Cox) test); comparisons between two groups (unpaired two-tailed t-test with Welch’s correction); comparisons among multiple groups (one-way ANOVA with Dunnett’s or Tukey’s post-hoc test); comparisons across multiple groups and conditions (two-way ANOVA with Bonferroni post-hoc test). Significance levels: ns, not significant; *p < 0.05, **p < 0.01, ***p < 0.001, ****p < 0.0001.

## Acknowledgement

We acknowledge NUSMed Confocal Microscopy Unit for providing confocal microscopy facilities used in this study. Some of the results here are produced based on data generated by the TCGA Research Network: https://www.cancer.gov/tcga. The funding support is provided by National Medical Research Council Individual Research Grant MOH-000944-01.

## Author contributions

G.X.Y. and M.W.: conceptualization, methodology discussion, data analysis, writing manuscript; G.X.Y.: data collection and initial analysis; M.W.: Funding, resources, supervision, review and editing; P.J.C.: data evaluation, manuscript writing and editing.

## Data availability

No large datasets were generated during the current study. All method and data details related to the presented results are available from the corresponding author upon request.

## Ethics approvals

All lab studies were under the guidance and approval by the Institutional Biosafety Committee and the Institutional Animal Care and Use Committee.

