## Supplementary Figures for "Concurrent inhibition of ICMT and RAF/MEK suppresses RAC1^P29S^-driven MAPKi resistance in BRAF^V600E^ melanoma by regulating TAZ activity"

### Figure S1

**A** Skin Cutaneous Melanoma (TCGA, Firehose Legacy)

| | Disease Free (Months) | Mutation Type | Mutation Site | Copy Number Alterations | Isoform |
|---|---|---|---|---|---|
| 1 | 102.1 | Driver | P29L | Gain | |
| 2 | 56.34 | Driver | P29L | Gain | |
| 3 | 11.01 | Driver | P29L | Diploid | |
| 4 | 6.27 | Driver | P29L | Gain | |
| 5 | 64.75 | Driver | P29S | Diploid | |
| 6 | 25.89 | Driver | P29S | Gain | |
| 7 | 24.64 | Driver | P29S | Gain | |
| 8 | 21.78 | Driver | P29S | Diploid | |
| 9 | 19.02 | Driver | P29S | Gain | |
| 10 | 18.69 | Driver | P29S | Diploid | |
| 11 | 13.07 | Driver | P29S | Diploid | |
| 12 | 8.97 | Driver | P29S | Not Profiled | |
| 13 | 5.78 | Driver | P29S | Gain | |
| 14 | 1.97 | Driver | P29S | Not Profiled | |
| 15 | 3.58 | Driver | P34S | Gain | |
| 16 | 22.31 | Driver | N111K | Diploid | Rac1b |
| 17 | 122.63 | VUS | V14E | Gain | |
| 18 | 33.38 | VUS | E31D | Gain | |
| 19 | 12.06 | VUS | P106L | Amplification | Unknown |
| 20 | 3.91 | VUS | P159L | Gain | Rac1b |

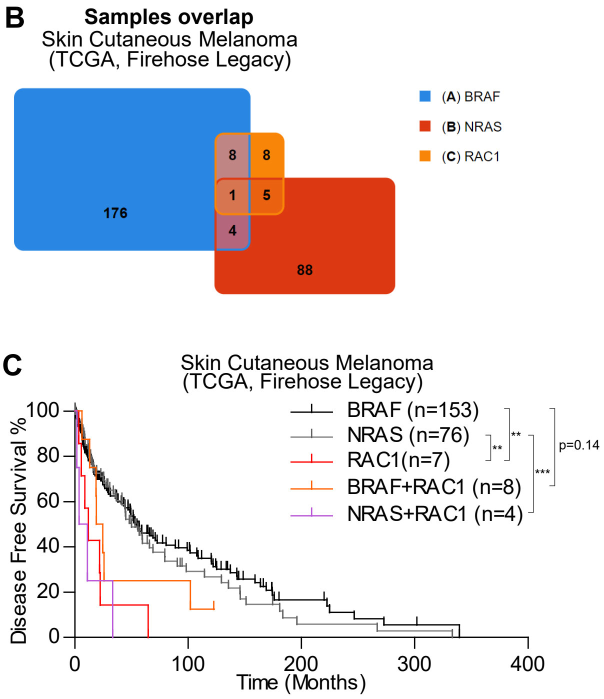


**Clinical significance of RAC1 alterations.**

**(A)** RAC1 alteration types for samples (n=20) in Skin Cutaneous Melanoma (TCGA, Firehose Legacy) Database. VUS, variant of uncertain significance.
**(B)** Co-occurrence of RAC1 alterations with BRAF or NRAS alterations in Skin Cutaneous Melanoma (TCGA, Firehose Legacy) Database.
**(C)** Percentage disease free survival by month for samples in (B). Statistical significance was determined by log-rank (Mantel-Cox) test for each indicated pair of curves: **p < 0.01, ***p < 0.001.

#### Figure S2

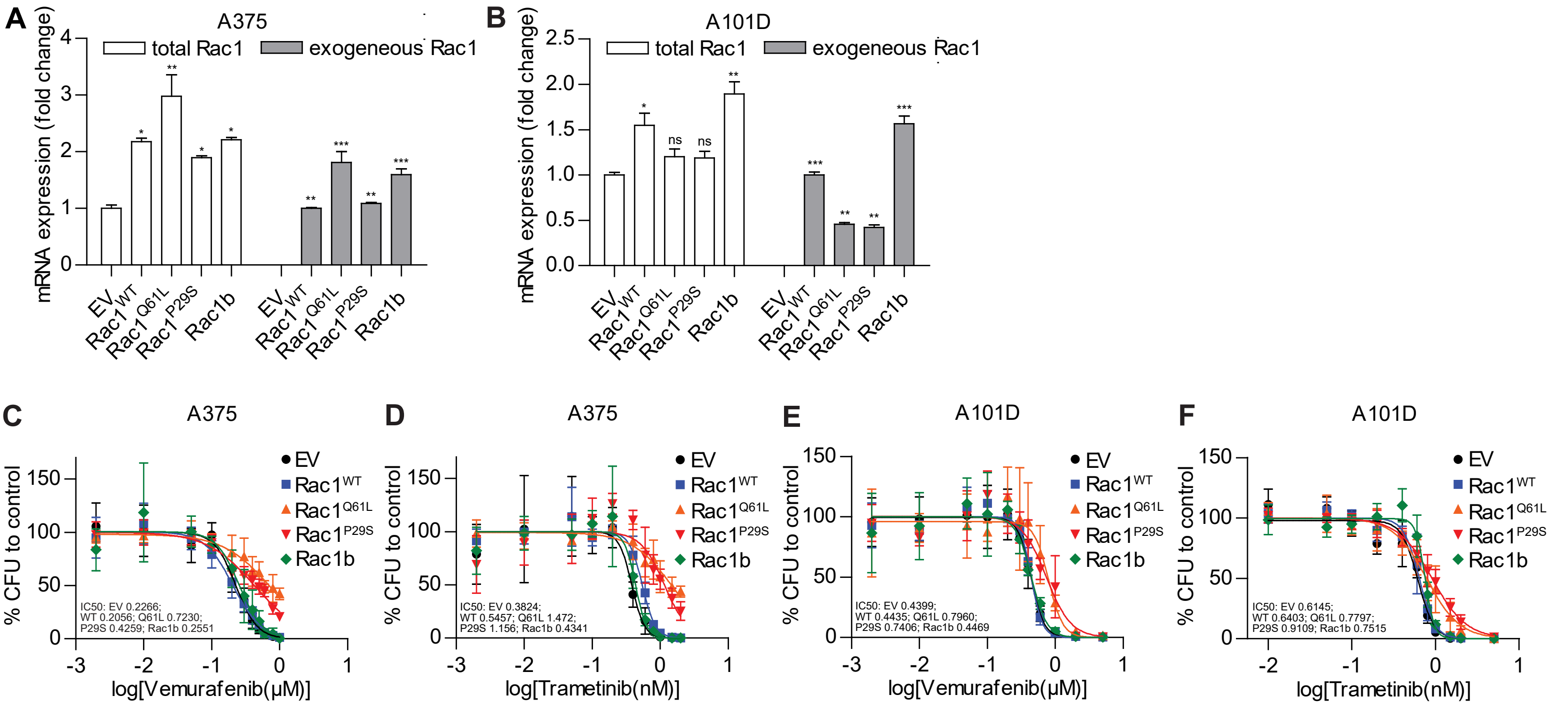


**Transgene expression and complete growth inhibition curves for soft agar colony formation in Figure 1.**

**(A)** Gene expression level of total RAC1 and exogeneous RAC1 in A375 cells transduced with EV, $RAC1^{WT}$, $RAC1^{Q61L}$, $RAC1^{P29S}$, or RAC1B, presented as mean ± S.E.M. Statistical significance compared with EV control was determined by one-way ANOVA with Dunnett's test: *$p < 0.05$, **$p < 0.01$, ***$p < 0.001$.
**(B)** Gene expression level of total RAC1 and exogeneous RAC1 in A101D cells transduced with EV, $RAC1^{WT}$, $RAC1^{Q61L}$, $RAC1^{P29S}$, or RAC1B, presented as mean ± S.E.M. Statistical significance compared with EV control was determined by one-way ANOVA with Dunnett's test: ns, not significant; *$p < 0.05$, **$p < 0.01$, ***$p < 0.001$.
**(C-D)** Colony inhibition curves of A375 cells expressing EV, $RAC1^{WT}$, $RAC1^{Q61L}$, $RAC1^{P29S}$, or RAC1B after 14-day incubation with (C) vemurafenib from 0.002μM to 1μM, and (D) trametinib from 0.002nM to 2nM.
**(E-F)** Colony inhibition curves of A101D cells expressing EV, $RAC1^{WT}$, $RAC1^{Q61L}$, $RAC1^{P29S}$, or RAC1B after 14-day incubation with (E) vemurafenib from 0.002μM to 5μM, and (F) trametinib from 0.01nM to 5nM.

#### Figure S3

A

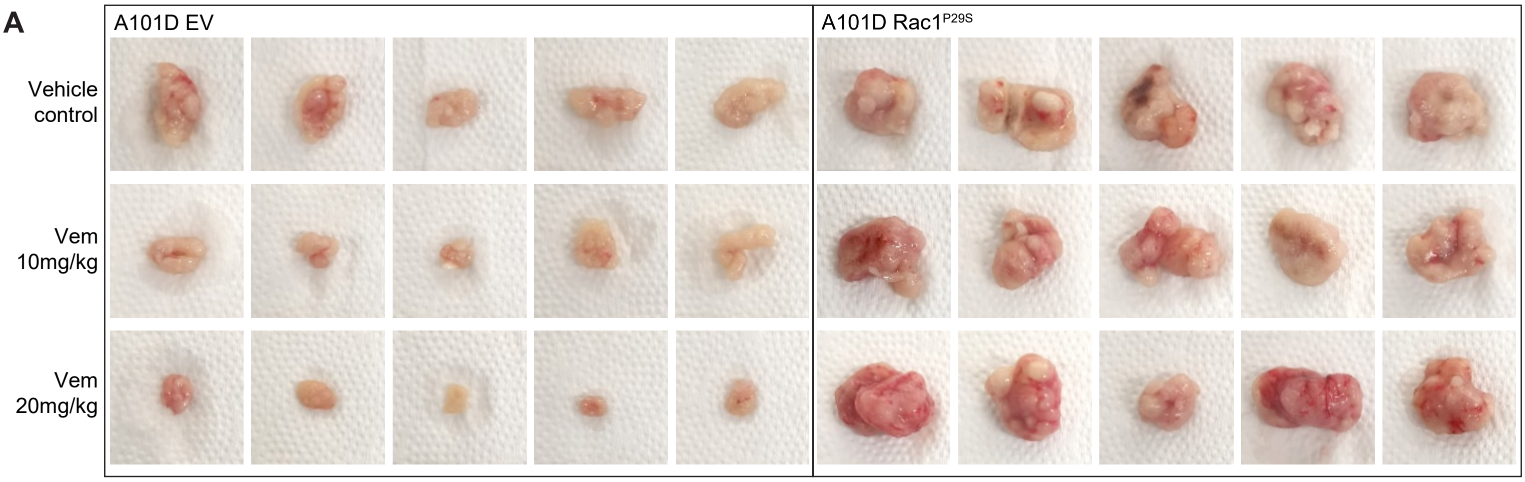


B

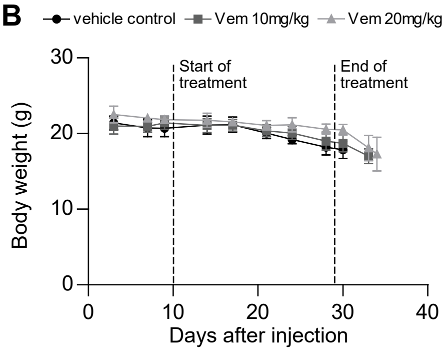


**Endpoint EV- or RAC1$^{P29S}$-transduced BRAF$^{V600E}$ melanoma xenografts under vemurafenib treatment and supporting data.**

**(A)** Complete images for tumors in Figure 1K.
**(B)** Body weight during treatment for mice of each group in Figure 1I.

### Figure S4

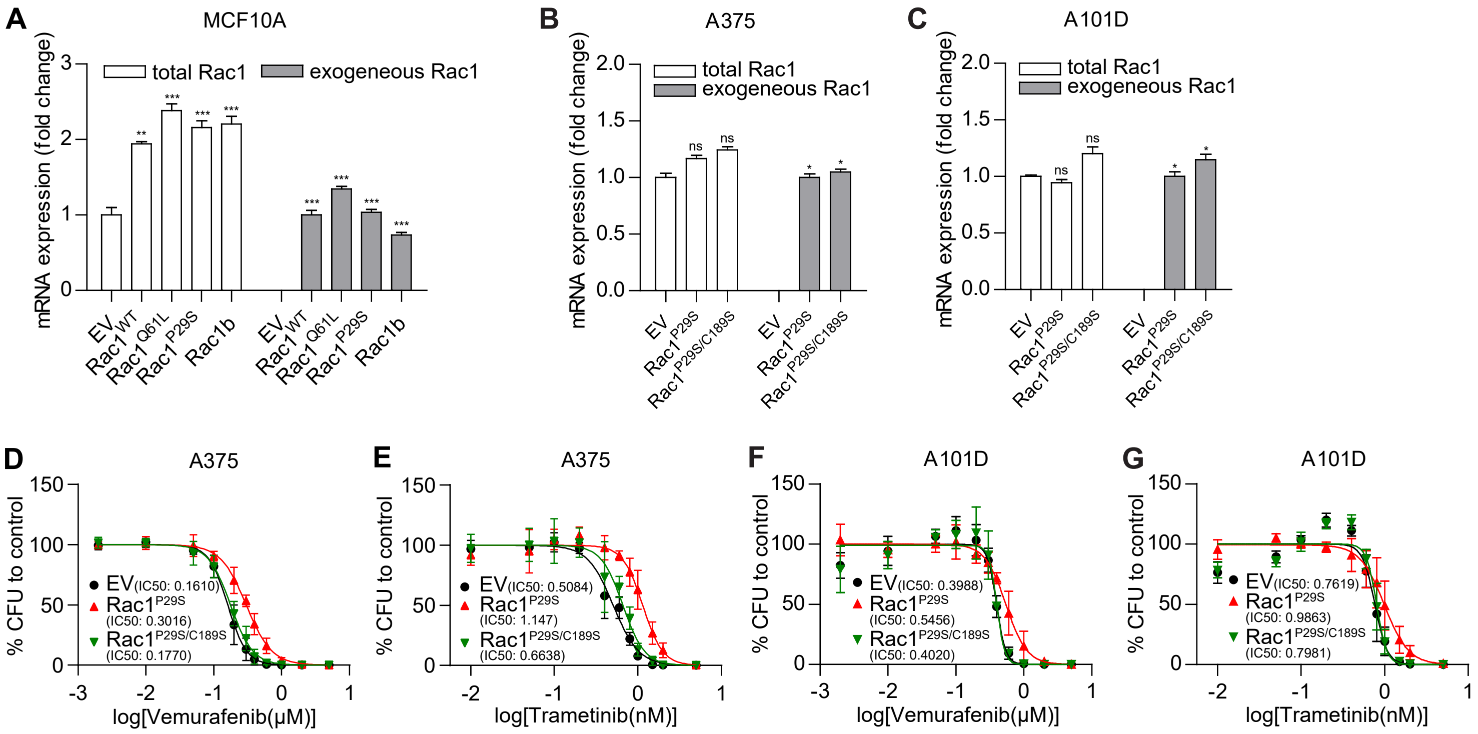


**Transgene expression and complete growth inhibition curves for soft agar colony formation in Figure 2.**

**(A)** Gene expression level of total RAC1 and exogeneous RAC1 in MCF10A cells transduced with EV, $RAC1^{WT}$, $RAC1^{Q61L}$, $RAC1^{P29S}$, or RAC1B, presented as mean ± S.E.M. Statistical significance compared with EV control was determined by one-way ANOVA with Dunnett's test: **$p < 0.01$, ***$p < 0.001$.
**(B)** Gene expression level of total RAC1 and exogeneous RAC1 in A375 cells transduced with EV, $RAC1^{P29S}$, or $RAC1^{P29S/C189S}$, presented as mean ± S.E.M. Statistical significance compared with EV control was determined by two-tailed unpaired t-test with Welch's correction: ns, not significant; *$p < 0.05$.
**(C)** Gene expression level of total RAC1 and exogeneous RAC1 in A101D cells transduced with EV, $RAC1^{P29S}$, or $RAC1^{P29S/C189S}$, presented as mean ± S.E.M. Statistical significance compared with EV control was determined by two-tailed unpaired t-test with Welch's correction: ns, not significant; *$p < 0.05$.
**(D-E)** Colony inhibition curves of A375 cells expressing EV, $RAC1^{P29S}$, or $RAC1^{P29S/C189S}$ after 14-day incubation with (D) vemurafenib from 0.002μM to 5μM, and (E) trametinib from 0.01nM to 5nM.
**(F-G)** Colony inhibition curves of A101D cells expressing EV, $RAC1^{P29S}$, or $RAC1^{P29S/C189S}$ after 14-day incubation with (F) vemurafenib from 0.002μM to 5μM, and (G) trametinib from 0.01nM to 5nM.

#### Figure S5

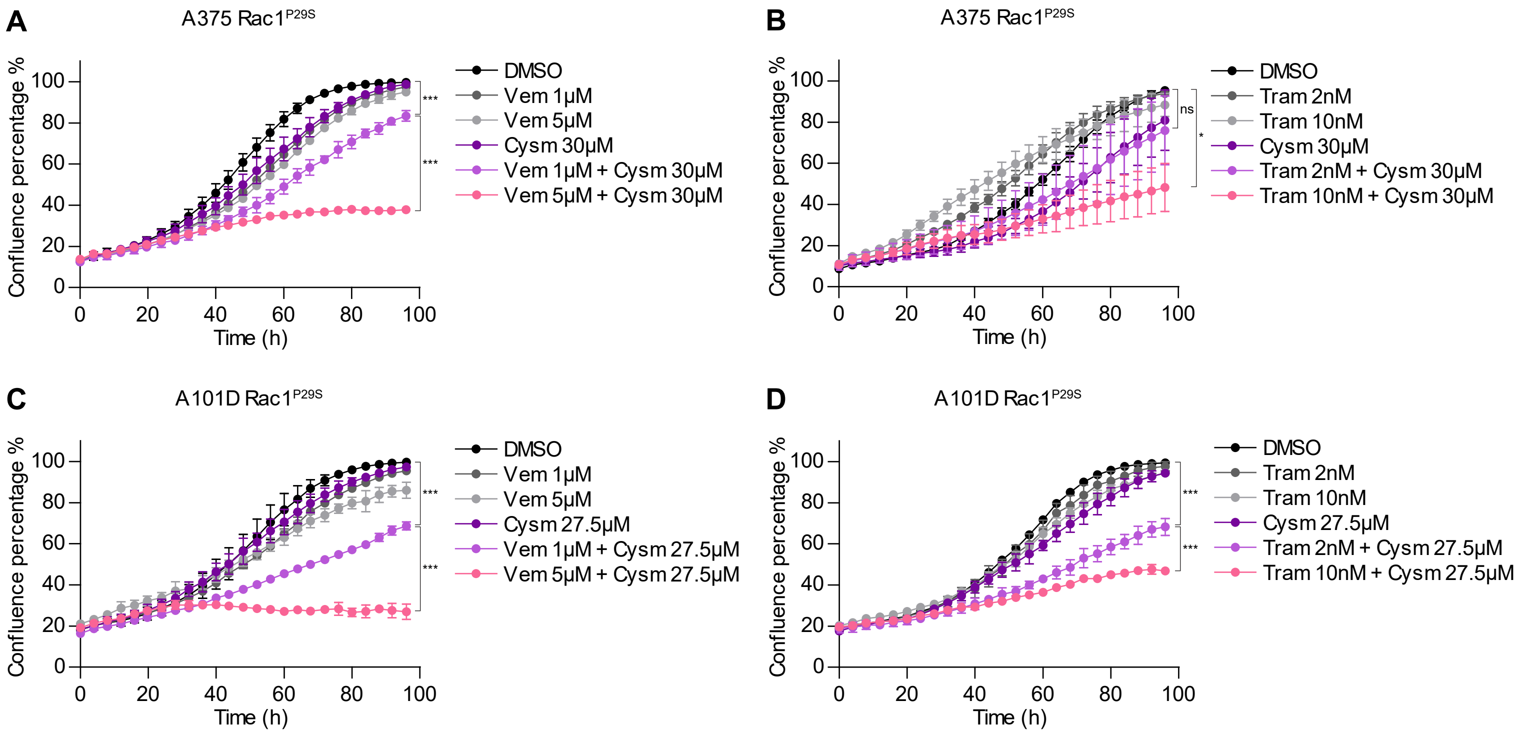


**ICMTi and MAPKi combination can elicit responses that MAPKi alone cannot induce.**

**(A-B)** Proliferation curves of RAC1$^{P29S}$-transduced A375 cells during 96h concurrent treatment of (A) 0, 1, 5μM vemurafenib plus 0, 30μM cysmethynil, and (B) 0, 2, 10nM trametinib plus 0, 30μM cysmethynil. Statistical significance was determined by one-way ANOVA with Tukey's test: ns, not significant; *p < 0.05, ***p < 0.001.

**(C-D)** Proliferation curves of RAC1$^{P29S}$-transduced A101D cells during 96h concurrent treatment of (C) 0, 1, 5μM vemurafenib plus 0, 27.5μM cysmethynil, and (D) 0, 2, 10nM trametinib plus 0, 27.5μM cysmethynil. Statistical significance was determined by one-way ANOVA with Tukey's test: ***p < 0.001.

#### Figure S6

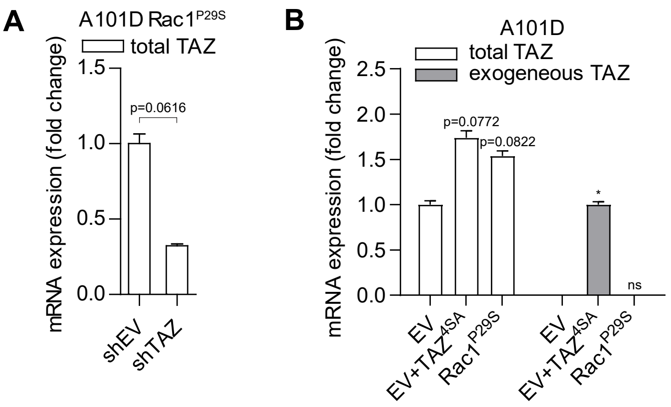


**TAZ expression levels in knockdown and over-expression study.**

**(A)** Analysis of shRNA knockdown efficiency in A101D RAC1$^{P29S}$ cells with qPCR, presented as mean ± S.E.M. Statistical significance was determined by two-tailed unpaired t-test with Welch's correction.

**(B)** Gene expression level of total TAZ and exogeneous TAZ in A101D EV, EV+TAZ$^{4SA}$ and RAC1$^{P29S}$ cells, presented as mean ± S.E.M. Statistical significance compared with EV control was determined by two-tailed unpaired t-test with Welch's correction: ns, not significant; $^{*}p < 0.05$.
